# Long-Term Performance of Utah Slanted Electrode Arrays and Intramuscular Electromyographic Leads Implanted Chronically in Human Arm Nerves and Muscles

**DOI:** 10.1101/2020.03.30.016683

**Authors:** Jacob A. George, David M. Page, Tyler S. Davis, Christopher C. Duncan, Douglas T. Hutchinson, Loren W. Rieth, Gregory A. Clark

## Abstract

**Objective:** We explore the long-term performance and stability of seven percutanous Utah Slanted Electrode Arrays (USEAs) and intramuscular recording leads (iEMGs) implanted chronically in the residual arm nerves and muscles of three human amputees as a means to permanently restore sensorimotor function after upper-limb.

**Approach:** We quantify the number of functional recording and functional stimulating electrodes over time. We also calculate the signal-to-noise ratio of USEA and iEMG recordings and quantify the stimulation amplitude necessary to evoke detectable sensory percepts. Furthermore, we quantify the consistency of the sensory modality, receptive field location, and receptive field size of USEA-evoked percepts.

**Main Results:** In the most recent subject, involving USEAs with technical improvements, neural recordings persisted for 502 days (entire implant duration) and the number of functional recording electrodes for one USEA increased over time. However, for six out of seven USEAs the number of functional recording electrodes decreased within the first two months after implantation. The signal-to-noise ratio of neural recordings and electromyographic recordings stayed relatively consistent over time. Sensory percepts were consistently evoked over the span of 14 months, were not significantly different in size, and highlighted the nerves’ fascicular organization. The percentage of percepts with consistent modality or consistent receptive field location between sessions (~1 month apart) varied between 0–86.2% and 9.1–100%, respectively. Stimulation thresholds and electrode impedances increased initially but then remained relatively stable over time.

**Significance:** This work demonstrates improved performance of USEAs, and provides a basis for comparing the longevity and stability of USEAs to that of other neural interfaces. Although USEAs provide a rich repertoire of neural recordings and sensory percepts, performance still generally declines over time. Future work should leverage the results presented here to further improve USEA design or to develop adaptive algorithms that can maintain a high level of performance.

## 1. Introduction

The Utah Slanted Electrode Array (USEA) consists of a 10 × 10 grid of electrodes with depths from 0.5 to 1.5 mm that penetrate peripheral nerves to enable single-unit neural recordings and intraneural microstimulation [1]. USEAs implanted in the peripheral nerves serve as a bi-directional sensorimotor neural interface, and have now been used to reanimate motionless limbs [2,3], to provide intuitive control of and sensory feedback from bionic arms [4–9], and to alleviate neuropathic pain [10,11]. The high electrode count and intrafasicular nature of the USEA yields improved specificity [6,12]. However, the long-term viability of chronically implanted USEAs has been uncertain [13].

The long-term viability of USEAs has become increasingly important as the field of neural prostheses moves rapidly towards clinical translation, with a current emphasis on multi-year, unsupervised, take-home trials [14–16]. Work with four prior amputee subjects demonstrated short-term (< 5 weeks) viability of USEAs as a sensorimotor interface for neural prostheses [7,8]. The present study extends these prior works by increasing the study duration well beyond four weeks and by providing additional data and performance analysis from iEMGs implanted in three new subjects. We quantify, for the first time, the long-term performance of USEA and iEMGs as a chronic sensorimotor neural interface for neural prostheses. We also highlight across-subject improvements in performance associated with technological advances in USEA design and showcase the longest duration of single-unit neural recordings from human peripheral nerve to date (502 days).

## 2. Methods

### 2.1 Human Subjects

We implanted USEAs and iEMGs into the residual arm nerves and muscles of three transradial amputees for recording and stimulation of neural signals and for recording muscle signals, respectively. The first amputee (subject 5; S5) was a 43-year-old male whose left and right forearms were amputated 24 years prior due to trauma [9,18]. The second amputee (subject 6; S6) was a 57-year-old male whose left foot and left forearm were amputated 13 years prior due to trauma [5,6,10,12]. The third amputee (subject 7; S7) was a 48-year-old male whose right forearm was electively amputated due to chronic Complex Regional Pain Syndrome with simultaneous intraoperative implantation of the USEA and iEMG devices [4,5].

All amputee participants were evaluated by a physician and psychologist for their willingness and ability to participate in the study. All surgeries and experiments were performed with informed consent from the participants and following protocols approved by the University of Utah Institutional Review Board and the Department of Navy Human Resources Proection Program.

### 2.2 Implanted Devices

We implanted two USEAs each in S5 and S6 (one in the median nerve and one in the ulnar nerve), and three USEAs in S7 (two in the median nerve and one in the ulnar nerve), as described in [5] (Fig. 1). For S7, one median-nerve USEA (S7-M1) was implanted more proximal and on the anterior side of the nerve under the dissected epineurium. The other median-nerve USEA (S7-M2) was implanted more distal and on the posterior side of the nerve under the dissected epineurium (i.e., the opposite side of the nerve relative to where the epineurium had been dissected open). After suturing back the dissected epineurium, the nerve was wrapped with an AxoGuard nerve wrap (AxoGen Inc., Alachua, FL, USA) to provide additional mechanical stability. A summary of the implanted devices can be found in Table 1.

**Figure 1.**
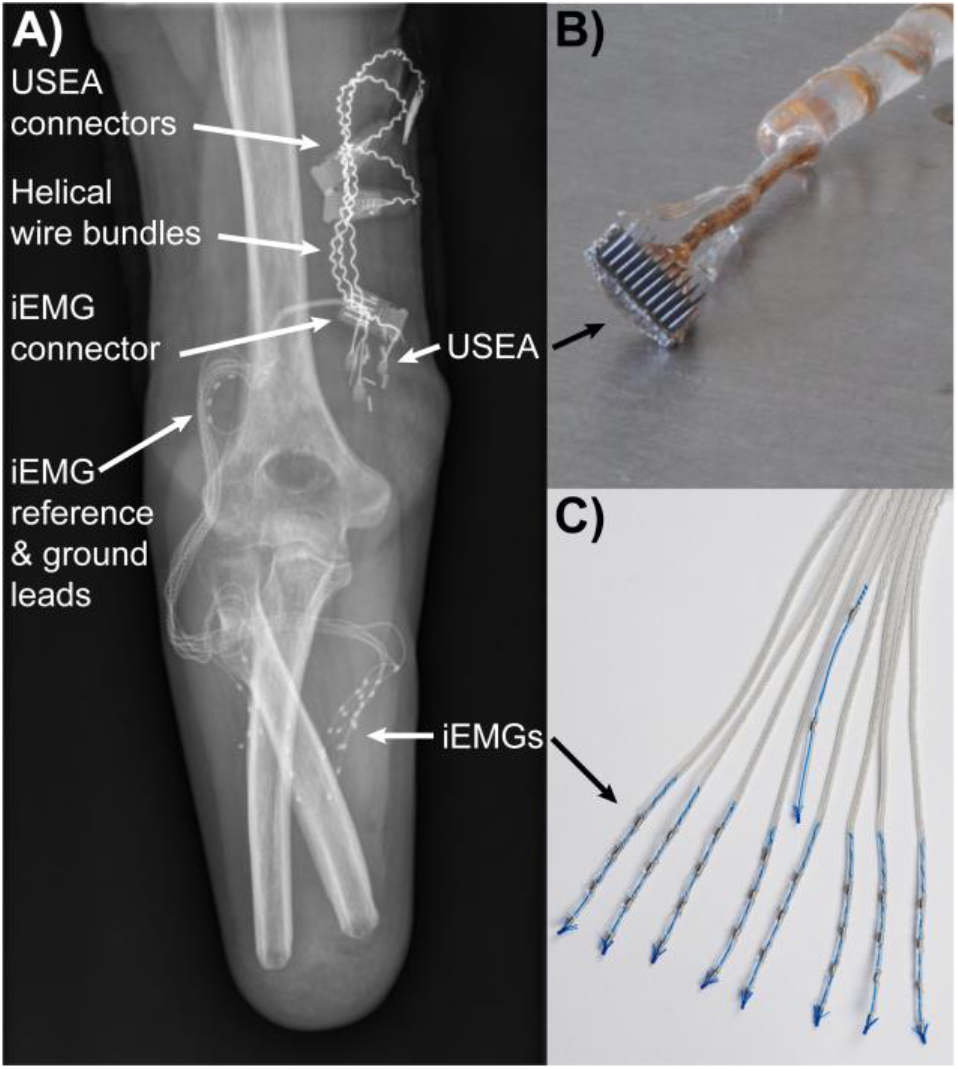
**A)** USEAs and iEMGs were implanted into three transradial amputees to restore sensorimotor function. **B)** USEAs were implanted into the residual median and ulnar nerves. **C)** iEMGs were implanted into the residual extrinsic hand muscles. Devices from S7 are shown. Images adapted from [5].

**Table 1:** Human Subjects and Implanted Devices

| Participant ID | Amputation | Study Duration | Device ID | Implanted Nerve |
| --- | --- | --- | --- | --- |
| S5 | Left transradial, 24 years prior | 84 days | S5-M | Median |
|  |  |  | S5-U | Ulnar |
| S6 | Left transradial, 13 years prior | 425 days | S6-M | Median |
|  |  |  | S6-U | Ulnar |
| S7 | Right transradial, during implant | 503 days | S7-M1 | Median |
|  |  |  | S7-M2 | Median |
|  |  |  | S7-U | Ulnar |

USEAs are silicon microelectrode arrays, with 100 electrode shafts, arranged in a 10 × 10 grid on a 4-mm × 4-mm base. Electrode shafts are spaced 400 μm apart, with shaft lengths varying from approximately 0.5–1.5 mm [1]. Four looped platinum wires were also implanted—two served as electrical ground and stimulation return, and two served as reference wires for recording.

All USEA tip metallization was performed by Blackrock Microsystems (Salt Lake City, UT, USA), using metallizations developed with support from the DARPA HAPTIX program. USEAs for S5 and S6 were tip-metalized with a pure platinum adhesion layer. USEAs for S7 were tip-metalized with a platinum silicide co-sputtered adhesion layer and utilized a more rapid annealing process. USEAs for S5 featured straight-connection wire bundles, whereas USEAs for S6 and S7 featured helical wire bundles. For S7, the helix was extended all way into the connector system, and additional silicone mechanical support was added to further reduce wire breakage [16].

All participants were also implanted with eight iEMG leads (with four contacts each) for a total of 32 electrode sites, with attempted targeting of each lead to different lower-arm extensor or flexor muscles. A separate iEMG lead (with two contacts) was implanted proximal and posterior to the elbow to provide an electrical reference and ground (Ripple Neuro LCC, Salt Lake City, UT, USA). Both USEA and iEMG electrodes exited the arm through a percutaneous incision and were mated with external Gator Connectors and associated Front Ends (Ripple Neuro LLC) for filtering and amplifying signals and for delivering stimulation [19].

USEAs and iEMGs were surgically removed after 84 days for participant S5 due an infection at the USEA percutaneous wire passage site. S5 recovered fully after USEA extraction and antibiotic treatment. USEAs and iEMGs were surgically removed after 425 and 503 days for participants S6 and S7, respectively, as the result of prior mutual agreements between the volunteer participants and the experimenters regarding study duration. Additional information regarding the surgical procedure can be found in [5].

### 2.3 Electrode Impedance

Electrode impedance was measured at 1 kHz and 10 nA with respect to the platinum ground wires using Trellis software (Ripple Neuro LLC). Impedances were measured at the start of each experimental session. We define functionally connected electrodes as electrodes with an impedance below 500 kΩ. The relatively high 500-kΩ threshold likely provides an overestimate of functionally connected electrodes for relatively low-impedance USEA electrodes, but has been utilized previously [7].

### 2.4 Recording Datasets

During each session, participants were instructed to perform three different sets of movements for ten seconds each, one after another, as described in [20]. These sets of movements were: 1) individual digit movements (isolated flexion and extension of D1–D5), 2) grasping (simultaneous flexion and extension of D1–D5), and 3) wrist movement (rotation, flexion/extension, and radial/ulnar deviation). These three sets of movements were performed one after another, and were then followed by a ten-second rest period where participants were instructed to relax and minimize their muscle activity. Only the middle eight seconds of each set of movements was analyzed to avoid transition delays due to participant reaction time. A total of 13, 65, and 46 datasets were collected for participants S5, S6, and S7, respectively. Additional experimental sessions were performed with the participants and have been reported previously [4–6,9,10,12].

### 2.5 Neural Recordings

Neural recordings were sampled at 30 kHz using the Grapevine Neural Interface Processor (Ripple Neuro LLC). Neural recordings were band-pass filtered with a 0.3 Hz first-order high-pass Butterworth filter and a 7500 Hz third-order low-pass Butterworth anti-aliasing filter. A digital high-pass filter with a cut-off frequency of 250 Hz (fourth-order Butterworth filter) was also applied to neural recordings.

Neural activity was detected by threshold crossings of an automated threshold set to negative five times the standard deviation of the electrode recordings during the rest period, as previously outlined in [8]. Threshold crossings were analyzed offline in MATLAB (MathWorks, Natick, MA, USA) to identify neural units. We defined neural units as a collection of threshold crossing events from a single electrode that have similar waveform shapes [21]. As noted in [21], neural units could be single-neuron action potentials, multi-neuron activity without well-isolated action potential shapes, or a collection of indistinct low-amplitude threshold crossings. For each threshold crossing, the voltage waveform was saved from the previous 100 μs to the subsequent 600 μs. Waveforms were designated as noise and removed from the data on the basis of visually non-overlapping outlier waveforms with interspike interval distributed around 60, 120 or 180 Hz.

Functional recording electrodes were defined as electrodes that have five times more threshold crossings during any of the three movement periods or rest period than the average number of threshold crossings across all electrodes during the rest period. Neural signal-to-noise ratio (SNR) was defined as the mean peak-to-peak value divided by two times the standard deviation during the rest period [8]. Because of the spike detection threshold (negative five times the standard deviation during rest), there is a lower limit of 2.5 for the SNR.

### 2.6 EMG Recordings

EMG recordings were sampled at 1 kHz using the Grapevine Neural Interface Processor. EMG recordings were band-pass filtered with cut-off frequencies of 15 Hz (sixth-order high-pass Butterworth filter) and 375 Hz (second-order low-pass Butterworth filter). Notch filters were also applied at 60, 120, and 180 Hz. EMG SNR was calculated in decibels as twenty times the root mean square of the EMG recordings during the three movement periods divided by the root mean square of the EMG during the rest period.

### 2.7 Neural Stimulation

Intraneural microstimulation was delivered via USEAs using the Grapevine Neural Interface Processor. All stimulation was delivered as biphasic, charge-balanced, cathodic-first pulses, with 200-μs phase durations, and a 100-μs interphase duration. Stimulation was delivered through individual USEA electrodes one by one. Stimulation was delivered in a pulsed fashion, with a 500-ms train of 100-Hz stimulation being delivered every second. Stimulation amplitude varied between 1–100 μA.

### 2.8 USEA-Evoked Percepts

For S5 and S6, we mapped the location, modality and detection threshold of the USEA-evoked percepts over time. Detection thresholds were identified by increasing stimulation current in 1–10-μA steps until a sensation was reliably detected. Participants reported the percept location by drawing the receptive field on an image of a hand. Receptive fields were then assigned to hand locations, corresponding with the five digits and anterior and posterior regions of the hand. The hand locations were based on the sensor locations of the LUKE Arm prosthesis (DEKA, Manchester, NH, USA), and included: Thumb, Index, Middle, Ring, Pinky, Median Palm, Ulnar Palm, and Back of Hand. Participants also selected the modality of the percepts from the following options: Vibration, Tightening, Tapping, Flutter, Pressure, Pain, Buzzing, Skin Stretch, or Proprioception (movement).

Mappings were completed at 15, 45, and 62 days post-implant for S5-M and at 20, 55, and 83 days post-implant for S5-U. For S6-M, mapping were completed at 11, 54, 83, 138, 173, 208, 242, 298, and 342 days post-implant. For S6-U, mappings were completed at 17, 55, 83, 138, 173, 208, 242, 298, and 354 days post-implant. Extensive mappings were not collective with S7 due to a narrow window of current between percept detection and discomfort (potentially due to prior Complex Regional Pain Syndrome) [4].

### 2.9 Percept Stability

We defined functional stimulating electrodes as electrodes that reliably evoked a detectable percept using stimulation currents below 100 μA. Electrodes that did not evoke a percept below 100 μA were not included in the stability analyses. To quantify stability of percept locations and percept modalities, we compared the percept location or modality for each electrode at adjacent timepoints. We define the percentage of stable electrodes between timepoints as the number of electrodes with consistent location or modality between the current timepoint and the previous timepoint out of the total number of electrodes that evoked sensations at both timepoints.

### 2.10 Statistical Analyses

Because data were not normally distributed, we performed a one-way non-parametric ANOVA (Kruskal-Wallis) for each USEA to determine if detection threshold or electrode impedance changed significantly between datasets (i.e., time as the sole factor and each USEA being an independent entity). If significance was found, we performed subsequent multiple comparisons (Wilcoxon rank-sum test) using adjusted alpha levels based on the Dunn-Sidak correction for multiple comparisons. Each individual comparison between timepoints was performed in an unpaired manner. Paired comparisons for electrodes were not performed because the number of overlapping electrodes between timepoints was much smaller than the total number of electrodes at each timepoint.

## 3. Results

### 3.1 The Number of Functionally Connected Electrodes Decreased Over Time for Two of Three Participants

The number of functionally connected electrodes (i.e., electrodes with an impedance under 500 kΩ) during the first sessions post-implant was 82 for S5-M, 71 for S5-U, 80 for S6-M, and 86 for S6-U, 82 for S7-U, 89 for S7-M1, and 56 for S7-M2. The number of functionally connected electrodes declined over time for USEAs implanted in S5 and S6. However, for S7, the number of functionally connected electrodes remained relatively constant for S7-U and S7-M1, and apparently increased over time for S7-M2 (Fig. 2A), perhaps due to a prior temporary faulty external connection or an increase in capacitive coupling or shunting. This improvement in the number of functionally connected electrodes across subjects is likely the result of the extended helical wire bundle and additional silicone mechanical support associated with the USEAs implanted in S7 [19], which presumably reduced lead-wire breakage and delimination at the bond pads of the USEA.

**Figure 2.**
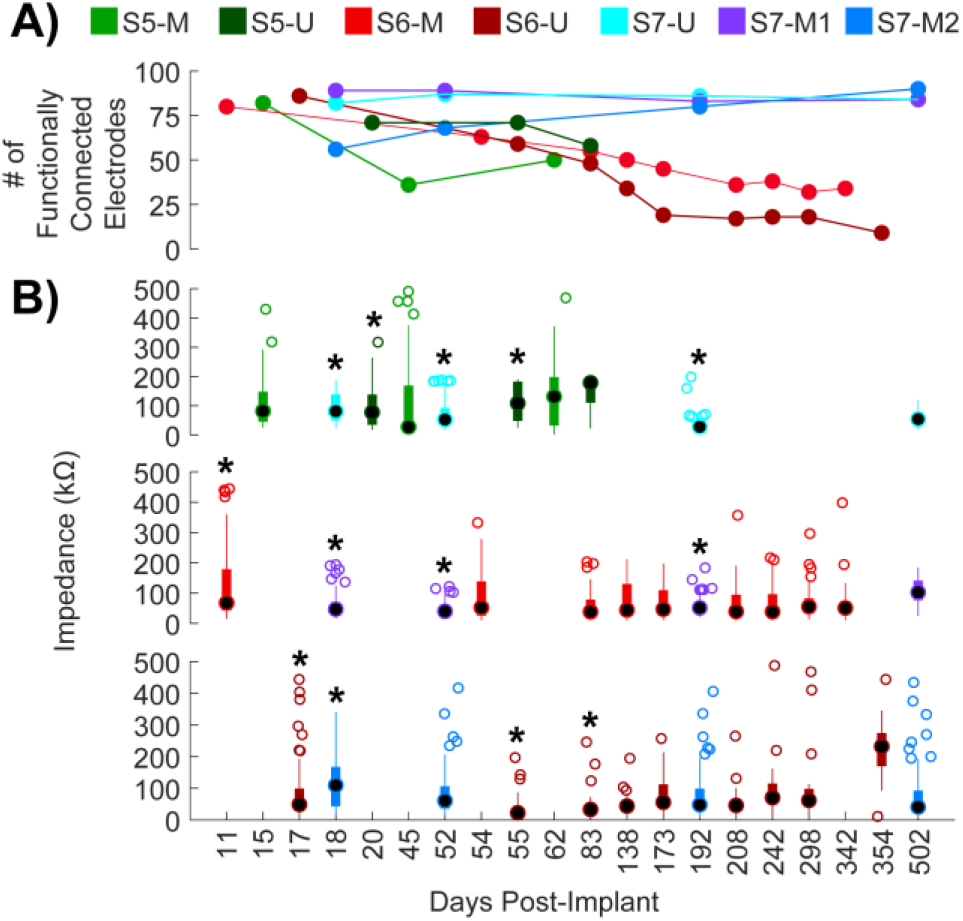
Impedance of functionally connected USEA electrodes over time. Timepoints are shown for each experimental session (ordinal scale). **A)** Functionally connected electrodes are defined as electrodes with an impedance below 500 kΩ. The number of functionally connected electrodes declined over time for USEAs implanted in S5 and S6, but maintained relatively constant for USEAs implanted in S7. **B)** Impedances of functionally connected electrodes were relatively stable over time. Black circles show median, thicker colored bars show IQR, thinner colored bars extend to the most extreme non-outlier values and open circles show outliers more than 1.5 times the IQR. * denotes statistically significant difference relative to a subsequent timepoint (*p* < 0.05, multiple comparisons with correction). **S5-M:** no significant differences. **S5-U:** day 20 and day 55 significantly different from day 83. **S6-M:** day 11 significantly different from days 83, 208 and 242. **S6-U:** day 17 significantly different from day 55; day 55 significantly different from days 173, 242, 298, and 342; day 83 significantly different from days 242 and 342. **S7-U:** day 18 significantly different from days 52, 192, and 502; day 52 significantly different from day 192; day 192 significantly different from day 502. **S7-M1:** days 18, 52 and 192 significantly different from day 502. **S7-M2:** day 18 significantly different from days 192 and 502.

### 3.2 Electrode Impedance Increased Initially and then Remained Relatively Stable

Generally, there were significant increases in electrode impedance within the first month post-implant, and few significant differences at later timepoints (Fig. 2B). With the exception of S5-M, electrode impedances during the first session were significantly different from impedances at least one subsequent session (*p* < 0.05, unpaired Wilcoxon rank-sum test with correction for multiple comparisons). The impedance and location of functionally connected electrodes on each USEA are shown in Fig. S1 and Fig. S2.

### 3.3 The Number of Functional Recording Electrodes Decreased Over Time for Two of Three Participants

The number of functional recording electrodes (i.e., electrodes recording neural activity) for S5 decreased from an initial 16 (at 7 days post-implant) to 1 over the 83-day study duration (Fig. 3, top). For S6, the study duration was extended and the number of functional recording electrodes decreased from an initial 20 (at 7 days post-implant) to zero within 131 days post-implant of the 425-day study duration (Fig. 3, middle). However, for S7, the number of functional recording electrodes remained relatively constant over the 502-day study duration (Fig. 3, bottom). For both S5 and S6, the maximum number of functional recording electrodes (26 and 25, respectively) occurred on the second experimental session (12 and 13 days post-implant, respectively). In contrast, the maximum number of functional recording electrodes for S7 (34) occurred on the 18^th^ experimental session, 143 days post-implant.

**Figure 3.**
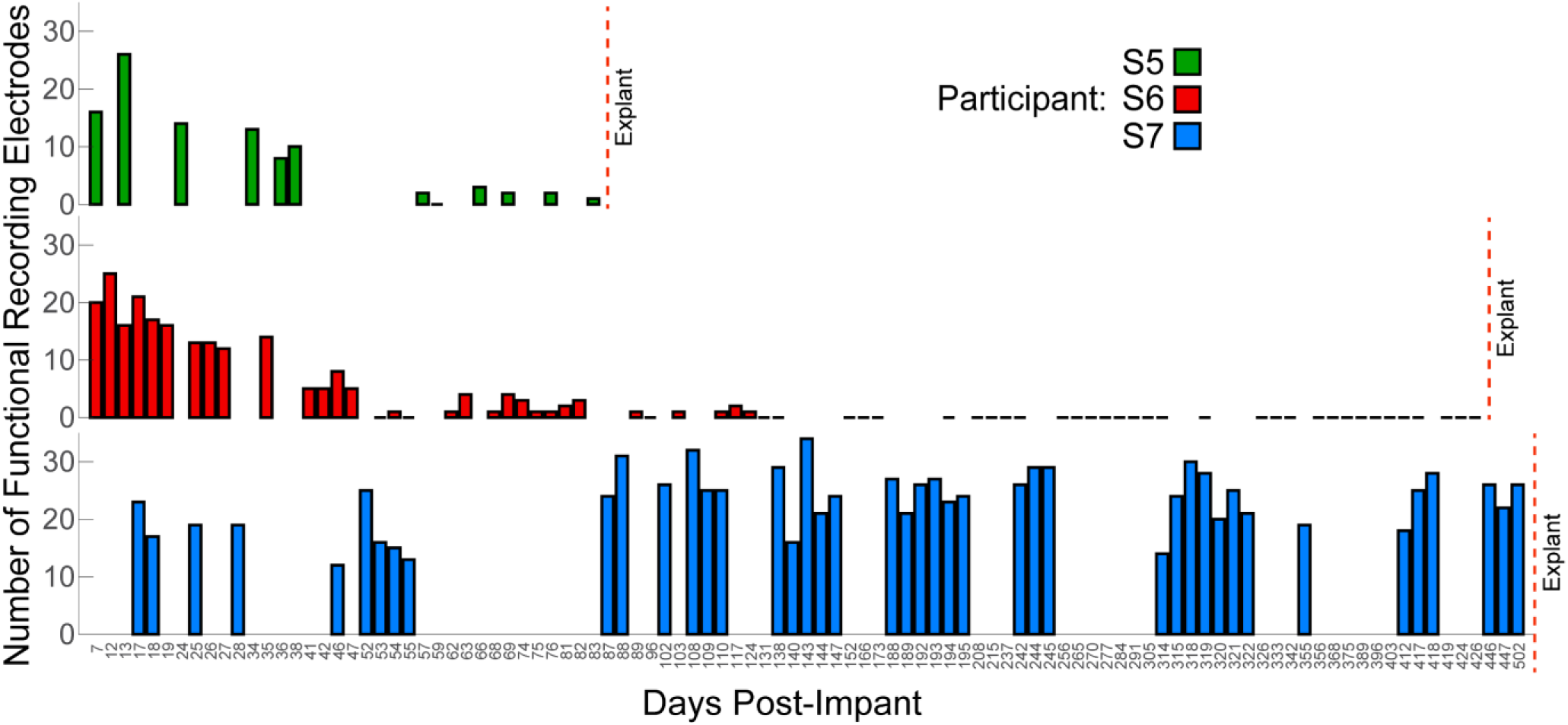
Number of functional recording electrodes over time. Timepoints are shown for each experimental session (ordinal scale). No measurements were taken on days with no bars. Zero functional recording electrodes were measured on days with flat black bars. For participants S5 (green) and S6 (red), the number of functional recording electrodes decreased overtime. For S7 (blue), the number of functional recording electrodes was relatively consistent over the duration of the implant.

All three USEAs implanted in S7 continued to function up to 502 days post-implant (Fig. 4). However, one USEA, S7-M2, contributed the majority of functional recording electrodes for S7. Interestingly, the number of functional recording electrodes for S7-M2 increased over the first two months (from an initial 13 at 18 days post-implant to 31 at 88 days post-implant). In constrast, the number decreased sharply within the first two months for all other USEAs in S7 and the USEAs in S5 and S6.

**Figure 4.**
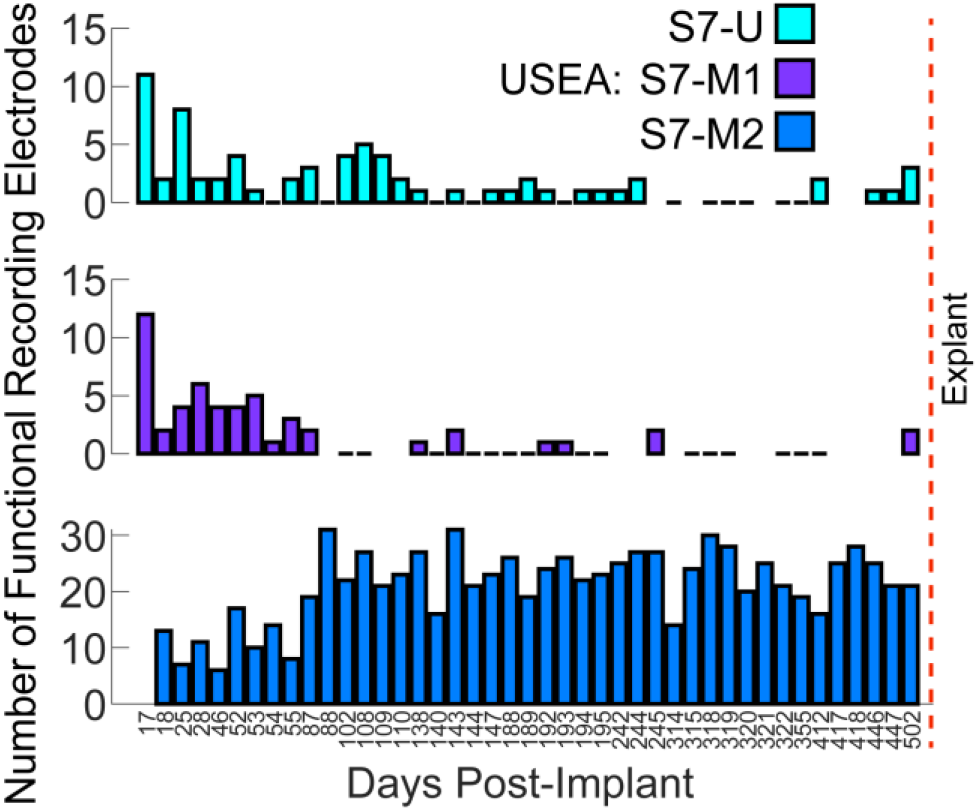
Number of functional recording electrodes for each USEA over time for participant S7. After the first two months, the vast majority of functional recording electrodes were located on one USEA (S7-M2, blue). Timepoints are shown for each experimental session (ordinal scale). No measurements were taken on days with no bars. Zero functional recording electrodes were measured on days with flat black bars.

The functional recording electrodes were distributed across the electrodes of the USEAs and corresponded to the location of functionally connected electrodes (Fig. 5). For S7-M2, a distinct diagonal band of functional recording electrodes was present (i.e., from ~E41 to ~E99) was was also consistent with the location of functionally connected electrodes (Fig. S2). Some of the descreases in the number of functional recording electrodes was likely due to the reductions in functionally connected electrodes due to issues such as lead wire breakage, rather than loss of recording ability per se.

**Figure 5.**
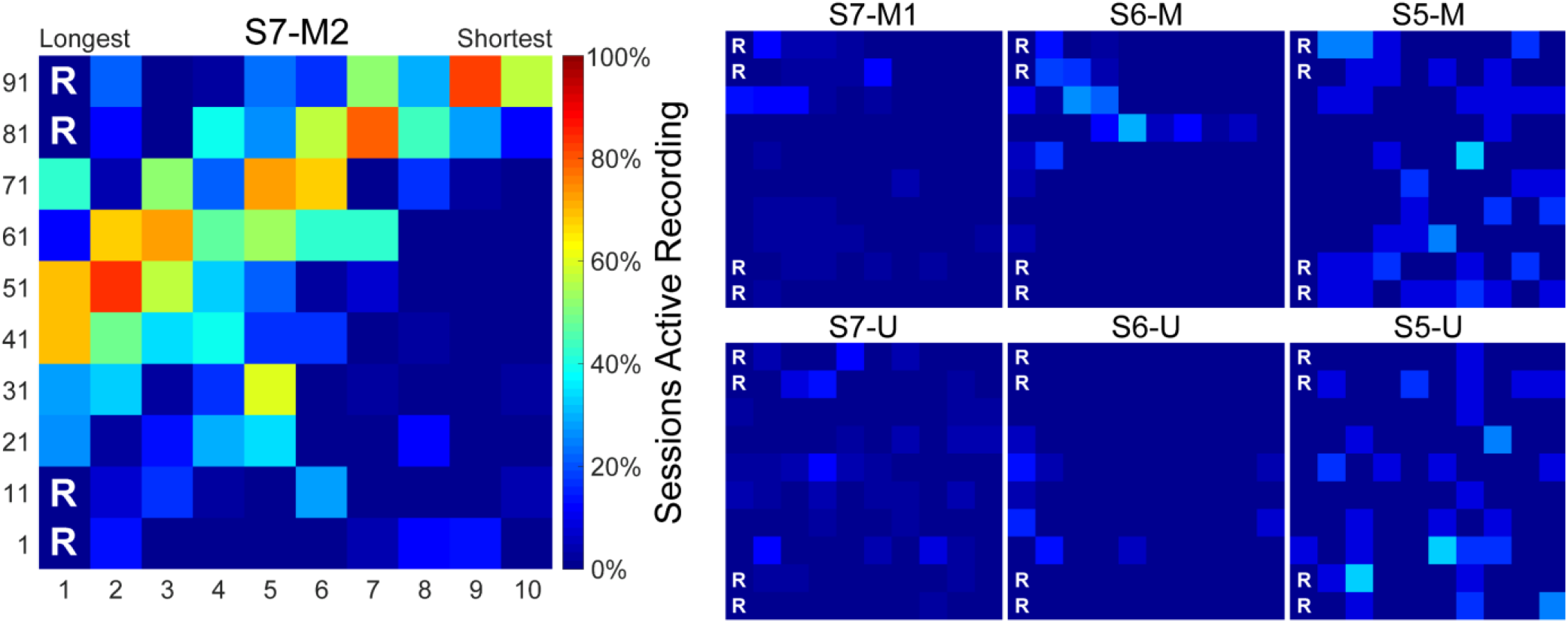
Percentage of sessions in which electrodes were functionally recording neural activity. Heatmaps show the 10 × 10 electrode grid on the USEA, with longest/deepest electrodes in columns 1 and shortest/shallowest electrodes in column 10. Electrodes 1, 11, 81, and 91 served as on-array references for recording. Hotter colors indicate a higher percentage of sessions in which that electrode was functionally recording neural activity.

### 3.4 Rich Repertoire of Neural and EMG Activity

Functional recording electrodes provided a rich repertoire of neural activity with dedicated subsets of units active primarily during unique sets of movements (Fig. 6). Some units were active primarily during finger movements (e.g., S7-M2, E75), others primarily during grasping (e.g., S7-M2, E64), and still others were primarily active during wrist movements (e.g., S7-U1, E55). Another subset of neural units were primarily active during rest (e.g., S7-M2, E33) while others were spotaneously active throughout the entire recording duration (e.g., S7-M2, E43). Importantly, individual neural units were not deterministically and uniquely related to a single movement; instead, most neural unit were still active to some degree across all three different types of movements.

**Figure 6.**
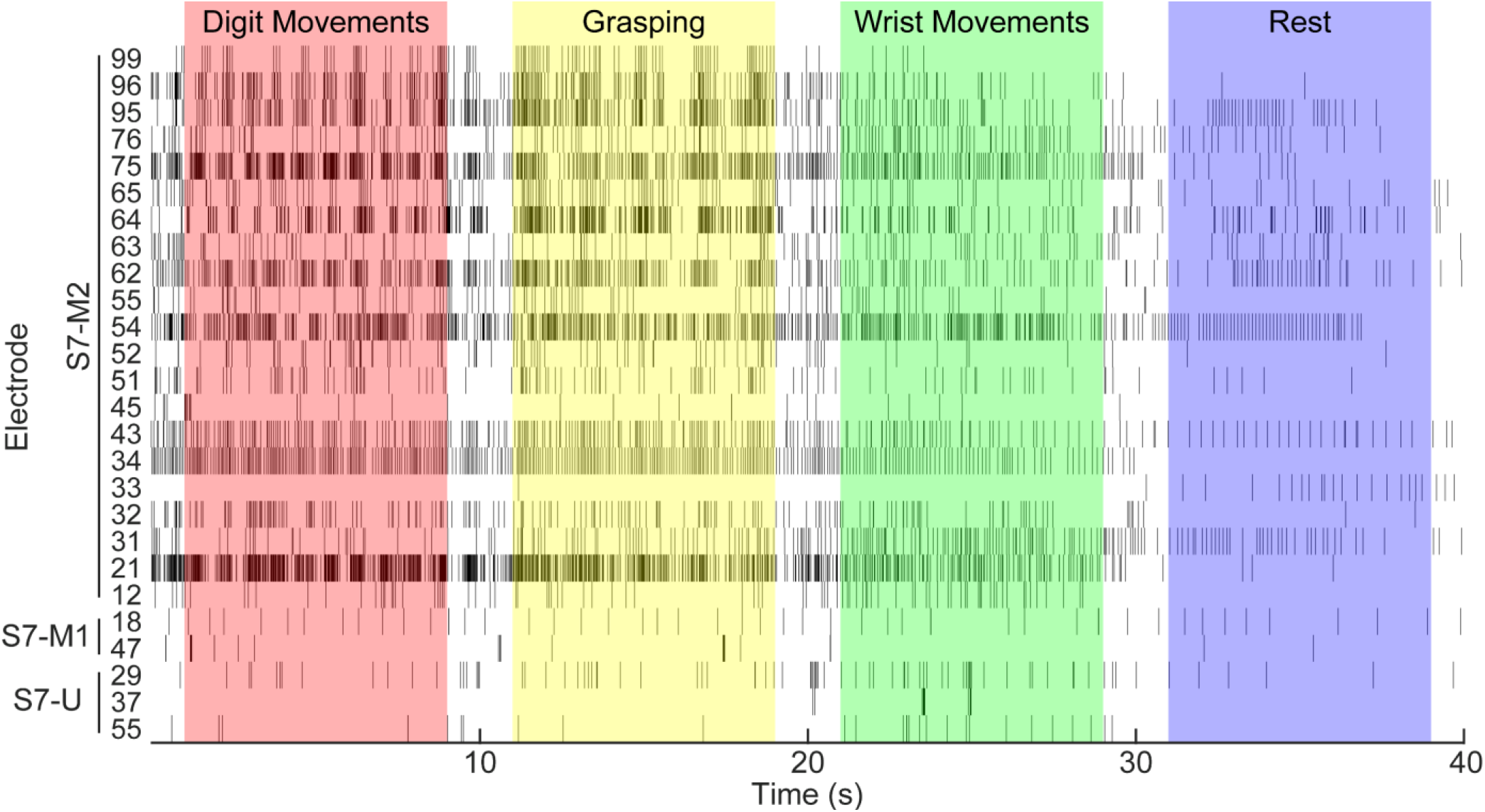
Neural units were successfully recorded long-term and were differentially recruited during different hand and wrist movements. Figure shows raster plot of neural units recorded from three USEAs (S7-M1, S7-M2, and S7-U) 502 days after they were implanted in participant S7. Vertical lines represent neural activity (i.e., threshold-crossing events). The participants were instructed to perform individual digit movements (~0–10 seconds, red), grasping (~10–20 seconds, yellow), and wrist movements (~20–30 seconds, green), followed by a rest period (~30–40 seconds, violet). Some units were active primarily during finger movements (e.g., S7-M2, E75), others primarily during grasping (e.g., S7-M2, E64), and still others were primarily active during wrist movements (e.g., S7-U1, E55). Another subset of neural units were primarily active during rest (e.g., S7-M2, E33) whereas others were spotaneously active throughout the entire recording duration (e.g., S7-M2, E43).

EMG activity recorded from the extrinsic hand muscles in the residual forearm provided a robust measure of motor intent, with distinct EMG ensembles for each different movement category (Fig. S3). EMG activity was still present for S7 even after long-term hand disuse and muscle atrophy. Abberent tonic activity was present on some channels – consistent with Complex Regional Pain Syndrome dystonia and myoclonus [22,23].

### 3.5 SNR of Neural and iEMG recordings

Neural units for participants S5–S7 had characteristic waveforms shapes, as shown previously for participants S1–S2 [8] (Fig. 7). The median (interquartile range (IQR)) SNR of neural recordings was 5.74 (1.75) for S5, 5.01 (1.54) for S6, and 5.57 (2.78) for S7. The median SNR tended to reside near the lower bound of SNR (2.5) based on the threshold-crossing detection criteria (Fig. 8). Neural units with large outlying SNRs (more than 1.5 times the IQR greater than the upper quartile) were likely recording EMG activity correlated with motor intent, as previously noted in [24]. The SNR and location of functional recording electrodes for each USEA are shown in Fig. S4.

**Figure 7.**
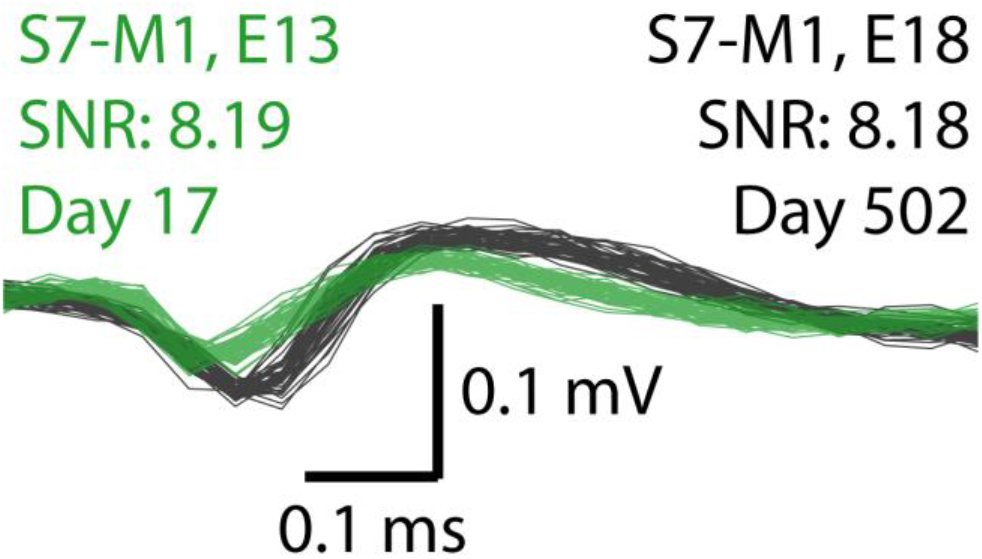
Representative (mean SNR) neural units recorded from S7-M1. Waveforms were recorded 17 days post-implant on electrode 13 (green) and 502 days post-implant on electrode 18 (black). Forty seconds of threshold-detected waveforms are overlaid for each electrode. Each waveform is 700 μs long, with 100 μs displayed prior to threshold crossing and 600 μs shown subsequent to threshold crossing.

**Figure 8.**
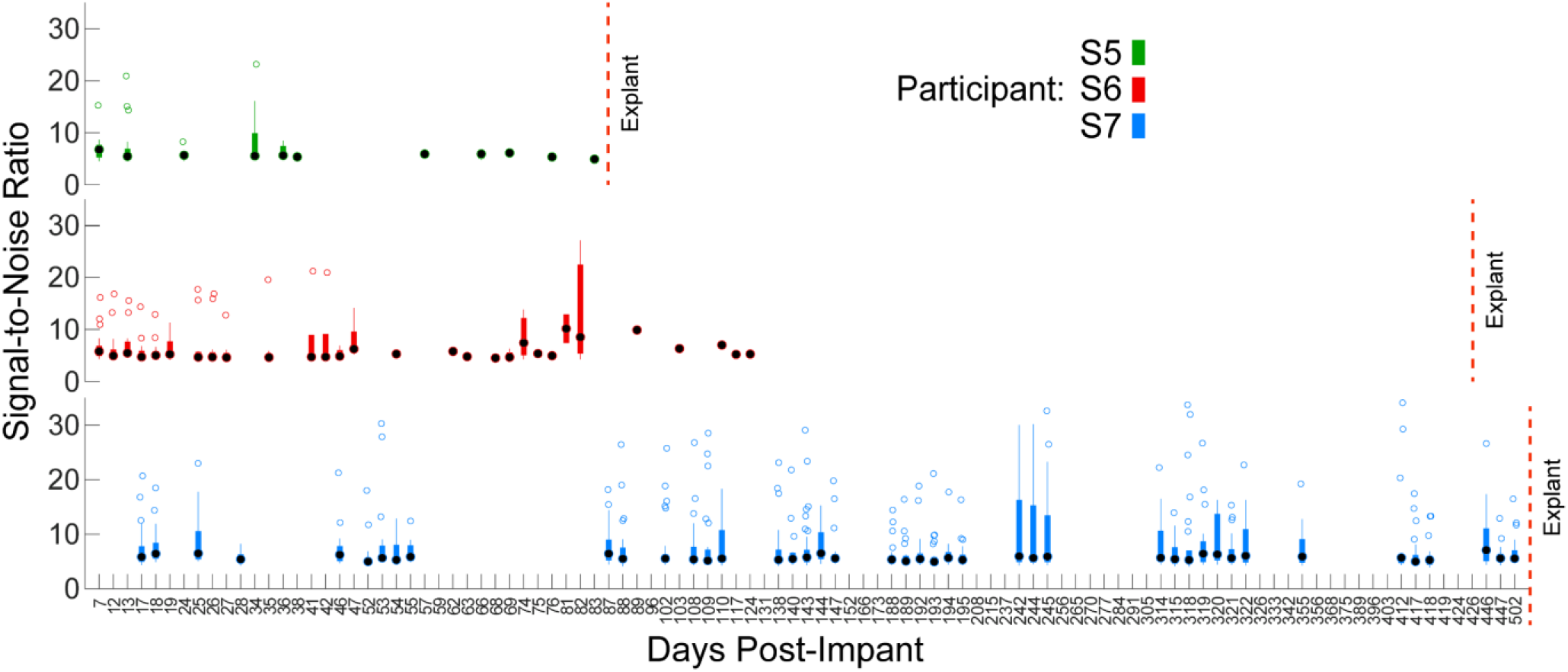
SNR of neural recordings over time. Timepoints are shown for each experimental session (ordinal scale). Black circles show median, thicker colored bars show IQR, thinner colored bars extend to the most extreme non-outlier values and open circles show outliers more than 1.5 times the IQR. Boxplots show the median SNRs for each functional recording electrode on that day, such that *N* is the number of functional recording electrodes on that day (values for *N* shown in Fig. 3). Because of the spike detection threshold (negative five times the standard deviation during rest), there is a lower limit of 2.5 for the SNR.

The median (IQR) SNR of EMG recordings was 45.54 (30.72) for S5, 48.02 (18.86) for S6, and 19.98 (19.49) for S7 (Fig. S5). EMG SNR for S7 was significantly lower than that of S5 and S6 (*p* < 0.05, unpaired Wilcoxon rank-sum test with correction for multiple comparisons), likely due to S7’s prior Complex Regional Pain Syndrome that resulted in multi-year hand disuse and muscle atrophy. Nevertheless, the SNR of iEMG recordings from these amputee participants was greater than the SNR of 32-channel surface EMG from intact participants using the same recording equipment (14.03 ± 4.43) [25]. iEMG recordings have also previously been reported to provide dexterous and intuitive control of bionic arms [5].

### 3.6 USEA Stimulation Evokes Numerous Cutaneous and Proprioceptive Percepts on the Phantom Hand

Intraneural microstimulation through individual USEA electrodes evoked localized sensations that were experienced on the phantom hand (Fig. 9). As reported previously [6], the location and modality of USEA-evoked percepts were consistent with known innveration patterns of the skin. A preponderance of percepts originated on the palmar side of the hand, and percepts were generally confined to the dermotome and myotome corresponding to the nerve in which the array was implanted. The reported modalities were also consistent with activation of cutaneous, proprioceptive and nociceptive nerve fibers. Cutaneous percepts were reported as “Pressure” (e.g., Merkel discs), “Tapping” or “Flutter” (e.g., Meissner corpuscles), “Vibration” (e.g., Meissner/Pacinian corpuscles), “Buzzing” (e.g., Pacinian corpuscles), and “Skin Stretch” (e.g., Ruffini endings). Proprioceptive percepts were reported as “Tightening” (e.g., Golgi tendon organs) and “Movement” (e.g., muscle spindles), and nociceptive percepts were reported as “Pain” (e.g., Aδ fibers). Infrequently, proprioceptive percepts were associated with motor fiber or h-reflex activation, as evidenced by visual muscle twitches on the residual limb. The median (IQR) percept sizes (percent of hand area) were 0.05% (0.67%) for S5-M, 2.72% (4.00%) for S5-U, 1.44% (4.14%) for S6-M, and 3.27% (6.68%) for S6-U (Fig. 10). These sizes are consistent with percepts evoked via longitudinal intrafascicular arrays [26]. There were no signfiicance differences in the size of USEA-evoked percepts over time (*p* = 0.65 for S5-M, *p* = 0.07 for S5-U, *p* = 0.44 for S6-M, *p* = 0.82 for S6-U; individual one-way Kruskal-Wallis).

**Figure 9.**
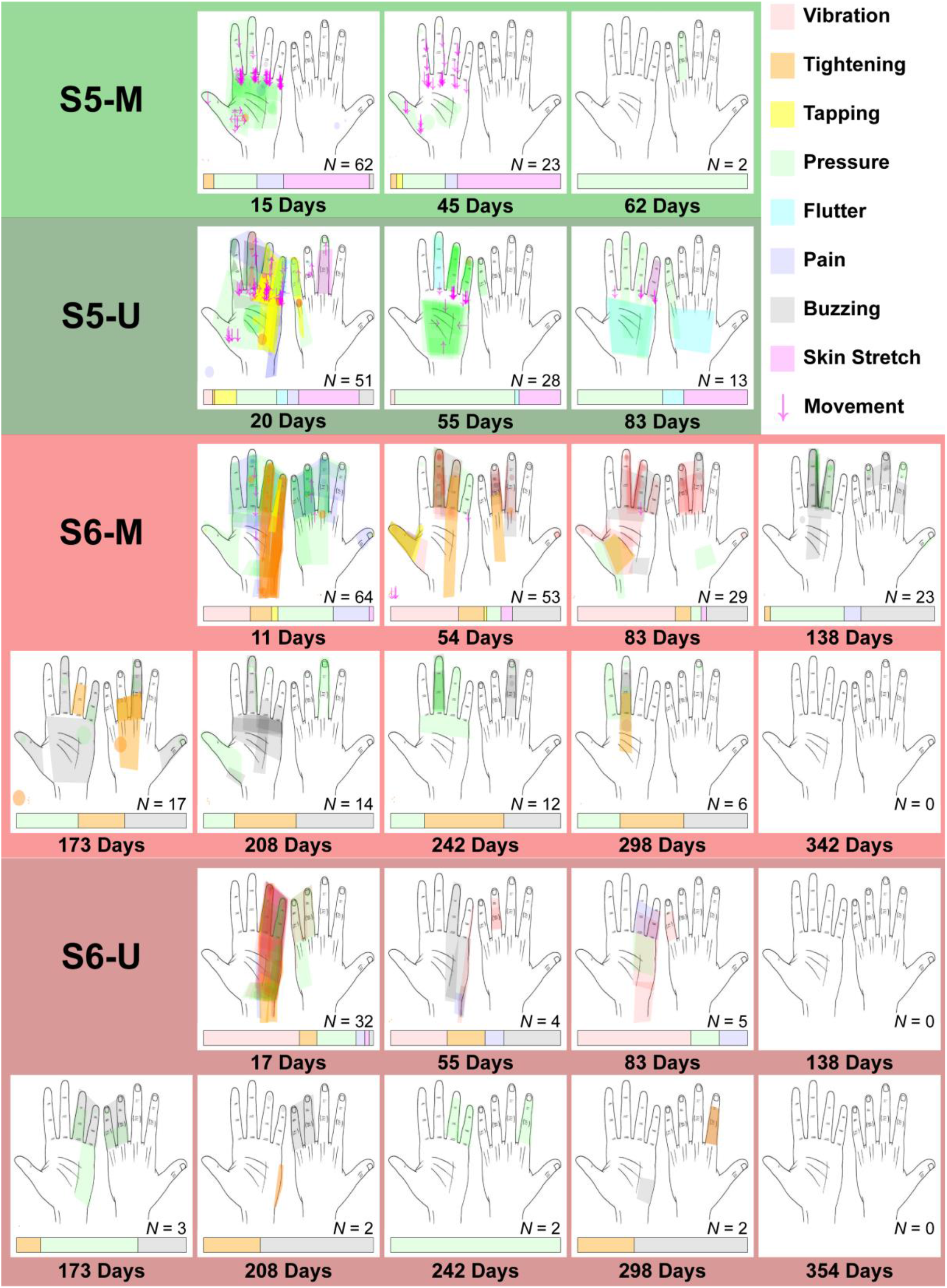
Receptive fields and modality distribution of USEA-evoked percepts over time. Percepts were evoked by stimulating each USEA electrode in isolation with increasing amplitude up to 100 μA. Participants self-reported the location of each percept by drawing the receptive field on an image of a left hand. Percepts marked off of the hand in the lower left (e.g., S5-U, 20 days post-implant) represent percepts experienced on the residual limb. Participants also self-reported the modality of each percept by selecting from nine possible modalities that corresponded to various cutaneous, proprioceptive or nociceptive sensory afferents. Each color drawn on the hand represents a different modality. Colors are transparent and overlaid such that more saturated colors represent multiple overlapping percepts with the same modality (e.g., brighter green on the palm versus faint green on D3, S5-U, 55 days post-implant). *N* shows the number of functional stimulating electrodes on that day and not necessary the total number of evoked percepts; a single functional stimulating electrode would occasionally evoke multiple percepts of different modalities. Horizontal bars below each hand image show the relative distribution of modality among the evoked percepts, such that larger horizontal space dedicated to a color represents a greater percentage of percepts with the modality associated with that color.

**Figure 10.**
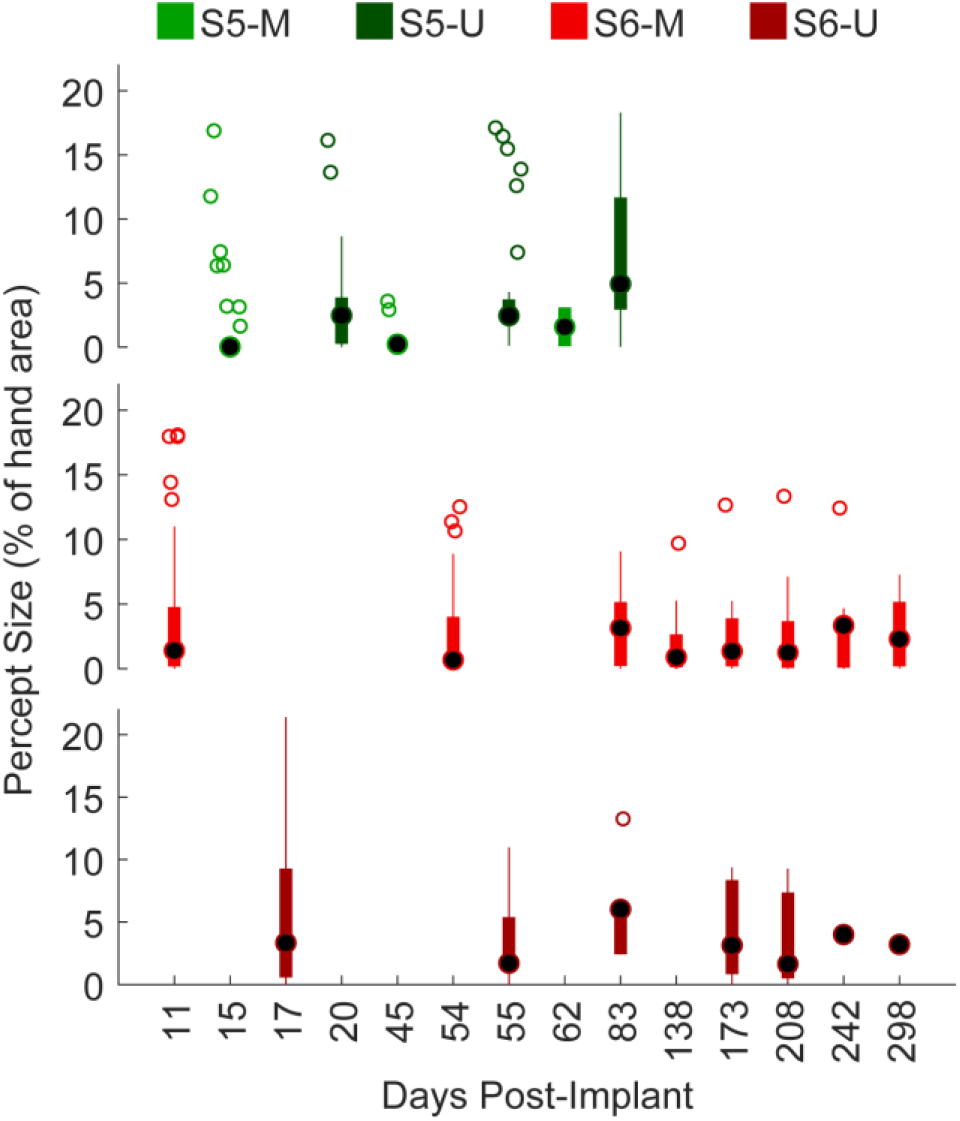
Size of USEA-evoked percepts over time. Timepoints are shown for each experimental session (ordinal scale). Size is represented as the percentage of hand area. Black circles show median, thicker colored bars show IQR, thinner colored bars extend to the most extreme non-outlier values and open circles show outliers more than 1.5 times the IQR. Four outliers greater than 22% (across all USEAs) are not shown for illustrative purposes. No significant differences were found across time for each USEA (*p* = 0.65 for S5-M, *p* = 0.07 for S5-U, *p* = 0.44 for S6-M, *p* = 0.82 for S6-U; individual one-way Kruskal-Wallis).

### 3.6 USEA-Evoked Percepts Reveal Fascicular Organization of Nerves

Adjacent USEA electrodes tended to evoke percepts with similar receptive fields – revealing the underlying fascicular organization of the nerve (Fig. 11). Electrode groupings based on receptive field were within 1-2 electrodes spacing (~400– 800 μm) and followed the orientation of the nerve (right to left on the USEA, proximal to distal on the nerve), cconsistent with the fascicular arrangement of human nerves [27]. For example, distinct fascicles for the thumb and middle finger were present on S6-M (Fig. 11, S6-M, row 2 in red and row 4 in yellow).

**Figure 11.**
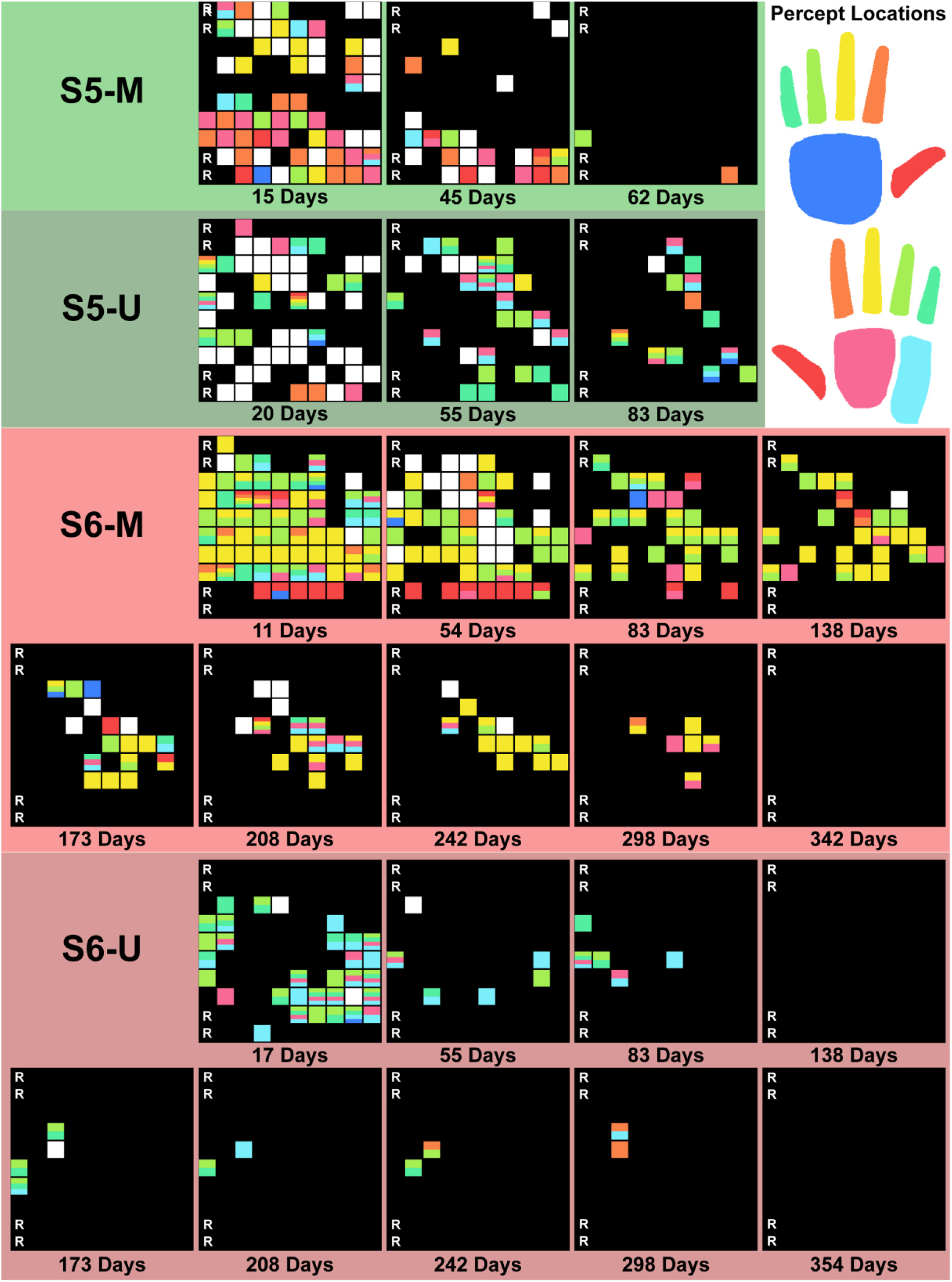
Fascicular arrangement of USEA-evoked percepts over time. Percepts were evoked by stimulating each USEA electrode in isolation with increasing amplitude up to 100 μA. Participants self-reported the location of each percept by drawing the receptive field on an image of a left hand. Percepts marked off of the hand (e.g., lower left, S5-U, 20 days post-implant) represent percepts experienced on the residual limb. Percepts were then mapped to discrete hand locations as defined by the colored hand regions in the top left. Images show the 10×10 electrode grid on the USEA, with longest/deepest electrodes in column 1 on the left and shortest/shallowest electrodes in column 10 on the right. Electrodes 1, 11, 81, and 91 served as on-array references for recording. The square for each electrode is colored to indicate the hand regions that the evoked percept spanned. Electrodes that evoked percepts spanning multiple hand regions are shown with multiple colors stacked vertically. Electrodes that evoked percepts that were on the residual limb or between discrete hand regions are represented by a white square. Fascicular orientation can be seen by distinct horizontal bands of color across the USEA (e.g., S6-M, red thumb region in row 2 and yellow middle finger region in row 4).

### 3.7 Location and Modality of USEA-Evoked Percepts Were Relatively Stable over Time

USEA-evoked percepts were relatively stable across a given pair of sessions (~1 month) with respect to modality and receptive field location (Fig. 12). For S6, a median of 73.9% of the percepts evoked via S6-M and a median of 100% of the percepts evoked via S6-U had consistent receptive fields between two adjacent mapping sessions. In contrast, for S5, a median of 9.1% of the percepts evoked via S5-M and a median of 39.6% of the percepts evoked via S5-U had consistent receptive fields between two adjacent mapping sessions.

**Figure 12.**
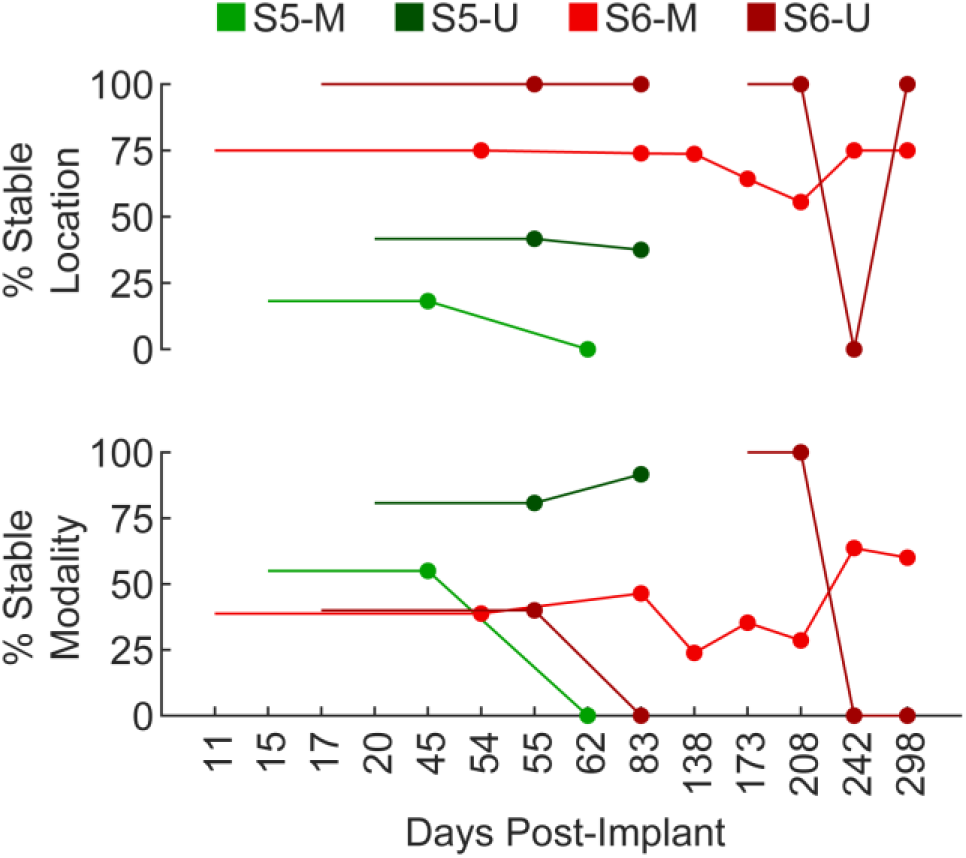
Stability of USEA-evoked percepts over time. Timepoints are shown for each experimental session (ordinal scale). Percent stable is defined as the number of electrodes with consistent location or modality between the current timepoint (dot) and the previous timepoint (line to the left) out of the total number of electrodes that evoked sensations at both timepoints. No percepts were evoked on S6-U between 83 days post-implant and 173 days post-implant.

The modality of percepts was generally less stable than the receptive field location. For S6, a median of 38.8% of the percepts evoked via S6-M and a median of 0% of the percepts evoked via S6-U had consistent modalities. In contrast, for S5, a median of 27.5% of the percepts evoked via S5-M and a median of 86.2% of the percepts evoked via S5-U had consistent modalities.

### 3.8 The Number of Functional Stimulating Electrodes Decreased Over Time

A maximum of 113 functional stimulating electrodes (63 from S5-M and 51 from S5-U) were reported for S5. A maximum of 96 functional stimulating electrodes (64 from S6-M and 32 from S6-U) were reported for S6. For both participants, the number of functional stimulating electrodes decreased over time (Fig. 13A). Only 15 functional stimulating electrodes were present for S5 prior to explantation, and zero functional stimulating electrodes were present for S6 prior to explantation. However, the loss of functional stimulating electrodes was not monotonic; zero functional stimulating electrodes were reported for S6-U at 138 days post-implant while four were reported at 173 days post-implant. Some of the descreases in the number of functional stimulating electrodes was likely due to the reductions in functionally connected electrodes due to issues such as lead wire breakage, rather than loss of stimulating ability per se.

**Figure 13.**
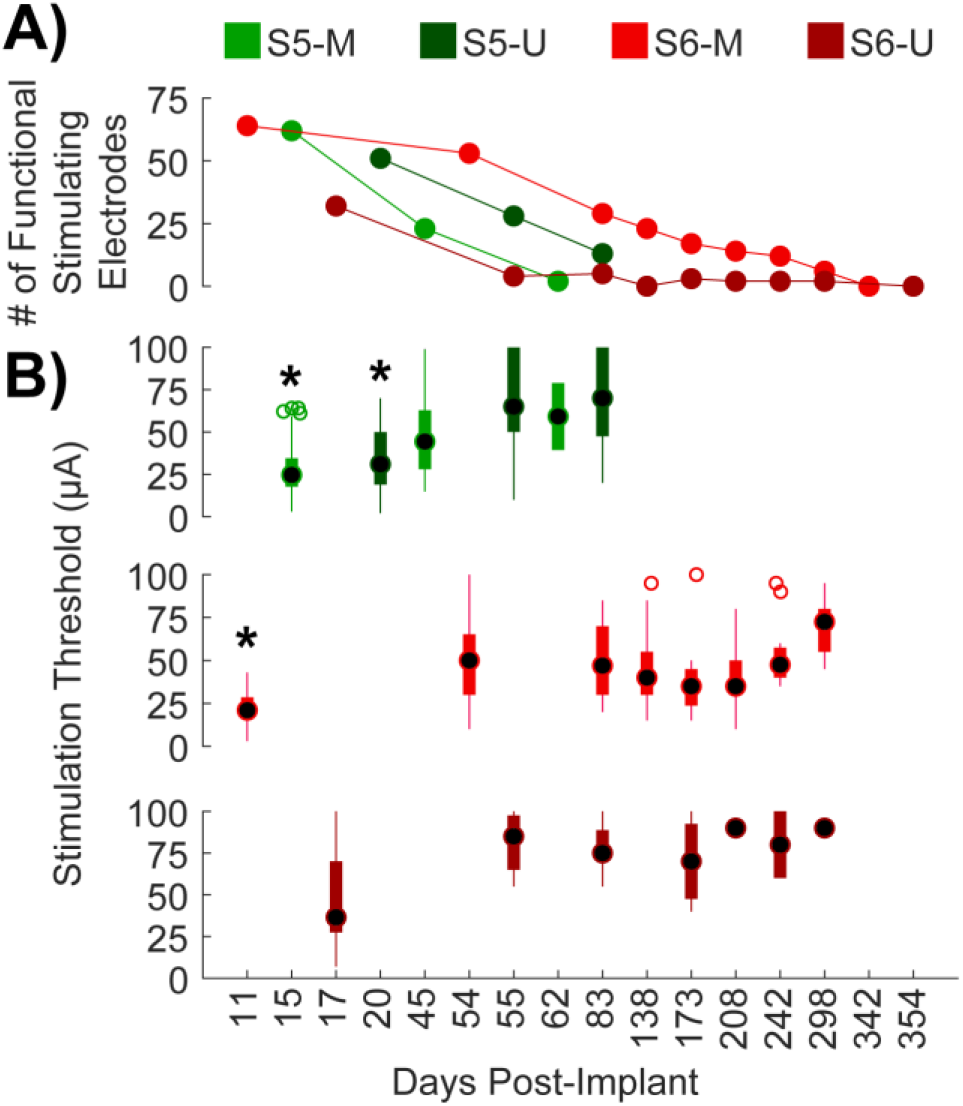
Number and detection threshold of functional stimulating USEA electrodes over time. Timepoints are shown for each experimental session and are not on a linear scale. **A)** The number of functional stimulating electrodes (i.e., electrodes that with a detection threshold below 100 μA) decreased over time. **B)** Detection thresholds increased over time. Detection thresholds for each electrode were identified by increasing the stimulation amplitude in 1 −10-μA step sizes until a percept was reliably detected. Black circles show median, thicker colored bars show IQR, thinner colored bars extend to the most extreme non-outlier values and open circles show outliers more than 1.5 times the IQR. Boxplots show the detection thresholds of functional stimulating electrodes, such that *N* = the number plotted in part A at the corresponding timepoint. * denotes statistically significant difference relative to a subsequent timepoint (*p* < 0.05, multiple comparisons with correction). **S5-M:** day 15 significantly different from day 45. **S5-U:** day 20 significantly different than days 55 and 83. **S6-M:** day 11 significantly different from days 54, 83, 138, 208, 242, and 298. **S6-U:** no significant differences.

### 3.9 Detection Thresholds Increased Initially and then Remained Relatively Stable

The median (IQR) detection thresholds during the first session were 25 μA (17 μA) for S5-M, 31 μA (31 μA) for S5-U, 21 μA (11 μA) for S6-M, and 36.5 μA (42.5 μA) for S6-U. In contrast, the detection thresholds during the last session with functional stimulating electrodes were 60 μA (40 μA) for S5-M, 70 μA (52.5 μA) for S5-U, 72.5 μA (25 μA) for S6-M, and 90 μA (0 μA) for S6-U (Fig. 13B) The thresholds during the first session and last session were only significantly different for S6-M (*p* < 0.05, unpaired Wilcoxon rank-sum test with correction for multiple comparisons). The lack of significance is likely attributed to the small sample sizes during the last session, the variability of detection thresholds within session, and the upper-limit of 100 μA. There was a significant increase from the first session to the second sessions for S5-M, S5-U, and S6-M (*p* < 0.05, unpaired Wilcoxon rank-sum test with correction for multiple comparisons). Functional stimulating electrodes were distributed across the electrodes of the USEAs and corresponded to the location of functionally connected electrodes (Fig. S6).

## 4. Discussion

USEAs have now been implanted in seven human amputees with increasingly longer study durations. This work leverages multiple months of data gathered from the most recent three human participants (S5–S7) to quantify the long-term performance of USEAs as a chronic sensorimotor neural interface.

The results presented here confirm previous findings that USEAs offer a unmatched ability to interface with numerous motor and sensory nerve fibers. USEAs analyzed here were able to record neural activity from up to 34 distinct electrodes and elicit up to 113 distinct sensory percepts through intraneural stimulation. Neural recordings have been shown to improve estimates of motor intent for bionic arms [5], and can be used to estimate motor intent when no residual muscles are available (e.g., from intrinsic hand muscles after amputation) [7]. Similarly, the ability to evoke numerous percepts corresponding to individual sensory afferents is critical for biomimetic sensory feedback for sensorized bionic arms [6]. USEAs are also capable of evoking multiple proprioceptive percepts; a difficult feat for other neural interfaces that have instead relied primarily on sensory substitution for proprioceptive feedback from bionic arms [28].

For S7, with the most recent version USEAs, a high level of functional recording electrodes was maintained throughout the 502-day study duration. The number of functional stimulating electrodes was not quantified over time for S7, but it is likely that functional recording electrodes are within close enough proximity to nerve fibers to also function as functional stimulating electrodes. It has also been shown that the amount of charge injected does not correlate with USEA electrode degradation [19].

In relative contrast, performance of USEAs dimished over time for S5 and S6 – shown most noteably through a decline in the number of functionally connected electrodes, functional recording electrodes and functional stimulating electrodes.

Multiple factors may have played a role in the differences in longevity between S5/S6 and S7. Likely of high importance, there were considerable technical improvements to the USEAs implanted in S7. Specifically, the helical wire bundle was extended all way into the connector system, and additional silicone mechanical support was added to reduce wire breakage [19]. There was noticeable delamination of the USEA backing in S5-M on explant, which likely occurred between 15 and 45 days post-implant and caused the upper half of the USEA to fail. S7 was also unique in that the devices were implanted during the amputation procedure, instead of over a decade after the amputation, but it is unclear how these biological differences would have interacted with USEA performance over time.

It is unlikely these technical improvements were the only contributor to the longevity of the devices, because a single USEA (out of three) in S7 provided the vast majority of functional recording electrodes. This USEA, S7-M2, was the only USEA to date to have been implanted on the posterior side of the nerve under the dissected epineurium (i.e., the opposite side of the nerve relative to where the epineurium had been dissected open). We speculate that this orientation may have increased mechanical stablity by allowing the residual epineurium and AxoGuard nerve wrap to provide more direct pressure holding the USEA into the nerve.

It’s also worth noting that the apparent loss of USEA electrodes was not monotonic. Both S7-M1 and S7-U had zero functional recording electrodes for up an extended period of time (up to 257 consectutive days) before recording neural units once again. Similarly, S6-U had zero functional stimulating electrodes at 138 days post-implant and four functional stimulating electrodes at 173 days post-implant. USEA performance seems to stabilize over time, although additional improvements are needed to allow the USEAs to consistently stabilize with a high level of functional electrodes, like was observed with S7-M2.

We defined functional recording electrodes as electrodes that recorded neural activity with threshold crossings five times the standard deviation of the noise. This threshold-crossing value has been previously used for USEAs and eliminates false-positive detection [7,8]. However, it’s possible that the USEAs in this study could have maintained a much higher level of functional recording electrodes with a lower threshold-crossing value. Similarly, we defined functional stimulating electrodes as electrodes that reliably evoked a detectable percept below 100 μA – a very conservative upper-limit for safety based on the Shannon limit [29]. Despite there being no functional stimulating electrodes on S6-M at 342 days post-implant, useful sensory percepts were still evoked with greater injected charge (e.g., using longer pulse widths, higher pulse amplitudes or simultaneous multi-electrode stimulation [6]). Hence, from a practical perspective, USEAs retained the functional ability to evoke sensory percepts, even when no individual electrode retained the ability to do so.

## 5. Conclusion

USEAs provide unmatched specificity as a bidirectional peripheral nerve interface. It is likely that recent improvements in USEA design have increased the long-term performance, but additional work is required to enable consistent longevity across all devices.

## Author Contributions

J.A.G. collected data for S6 and S7, analyzed data, and wrote the manuscript. D.M.P. collected data for S5. T.S.D. assisted with data collection for S5 and S6, and assisted with data analysis. C.C.D. provided clinical support. D.T.H. performed surgeries and provided clinical support. L.W.R. designed and developed USEAs. G.A.C. oversaw all methods and analyses and participated in experimental sessions. All authors contributed to the revision of the manuscript.

## Acknowledgements

We thank the amputee participants involved in this study for their willingness to contribute their lifes to improve science and the lives of future amputees. We acknowledge Dr. David Kluger, Dr. Suzanne Wendelken, Dr. Mark Brinton and Michael Paskett for assisting with participant scheduling and occassional data collection.

This work was sponsored by the Hand Proprioception and Touch Interfaces (HAPTIX) program administered by the Biological Technologies Office (BTO) of the Defense Advanced Research Projects Agency (DARPA) through the Space and Naval Warfare Systems Center, Contract No. N66001-15-C-4017. Additional sponsorship was provided by the National Science Foundation (NSF) through grant No. NSF ECCS-1533649, the NSF Graduate Research Fellowship Program Award No. 1747505, and the University of Utah Department of Veterans Affairs. The content herein represents the views of the authors, not their employers’ or funding sponsors’.

---

**Figure S1.**
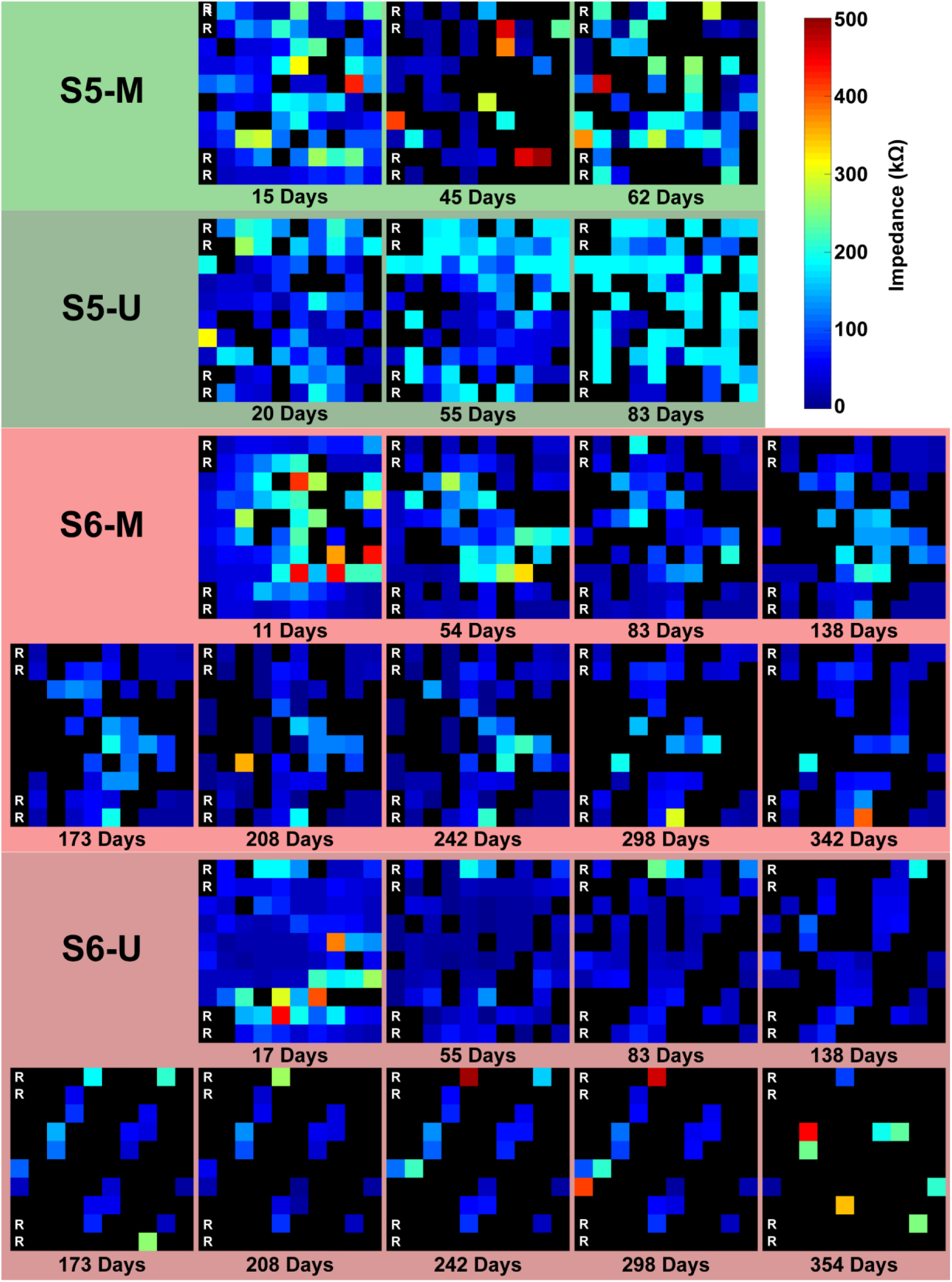
Impedance of USEA electrodes implanted in S5 and S6 over time. Heatmaps show the 10×10 electrode grid on the USEA, with longest/deepest electrodes in column 1 on the left and shortest/shallowest electrodes in column 10 on the right. Electrodes 1, 11, 81, and 91 served as on-array references for recording. Hotter colors indicate a higher impedance. Impedances above 500 kΩ are shown as black and are indicative of broken wires or faulty connections. Impedances of zero are show as the darkest blue and are indicative of faculty connections.

**Figure S2.**
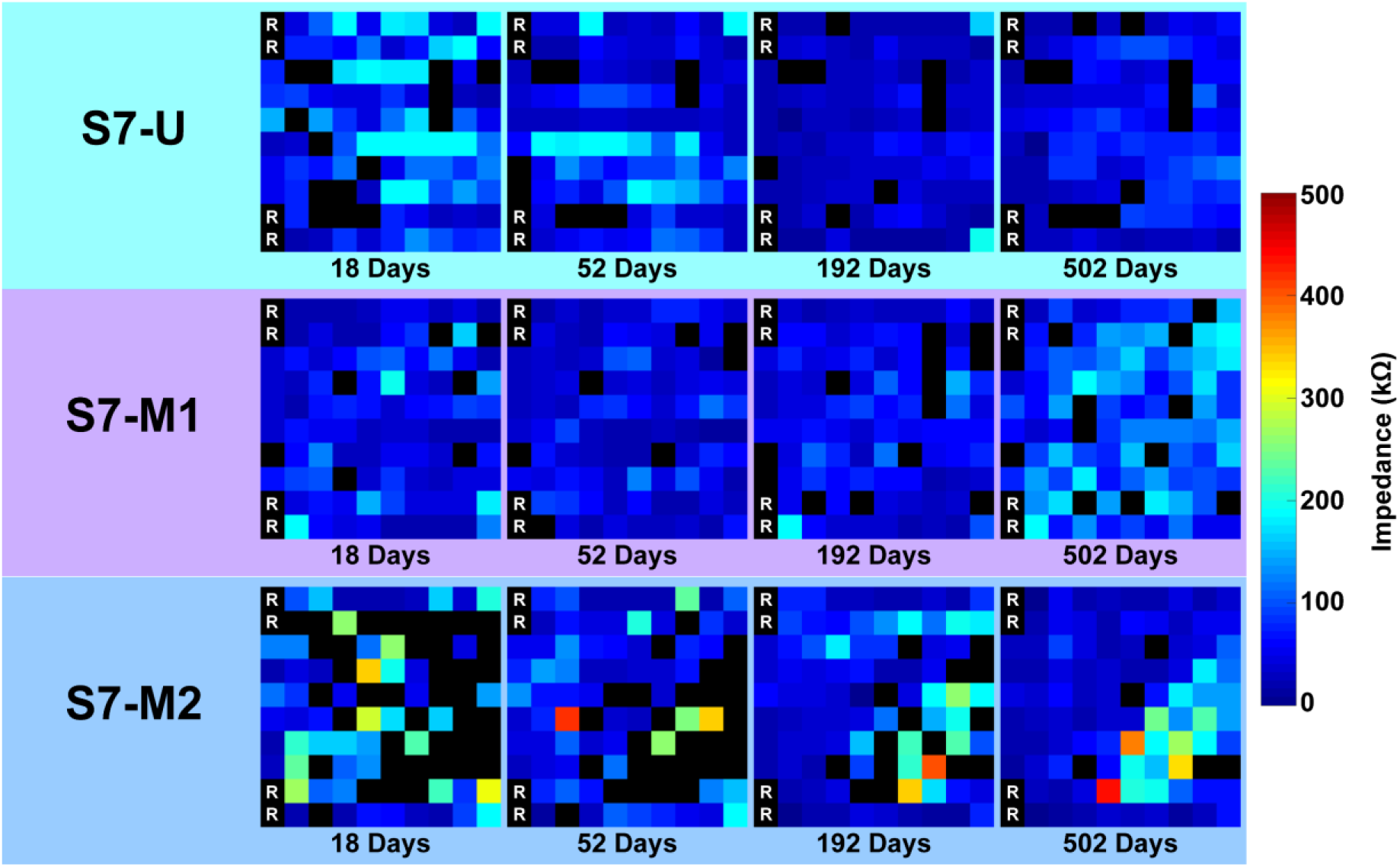
Impedance of USEA electrodes implanted in S7 over time. Heatmaps show the 10×10 electrode grid on the USEA, with longest/deepest electrodes in column 1 on the left and shortest/shallowest electrodes in column 10 on the right. Electrodes 1, 11, 81, and 91 served as on-array references for recording. Hotter colors indicate a higher impedance. Impedances above 500 kΩ are shown as black and are indicative of broken wires or faulty connections. Impedances of zero are show as the darkest blue and are indicative of faculty connections. Decreases in impedance down from 500 kΩ are likely the result of a prior temporary faulty external connection or an increase in capacitive coupling or shunting.

**Figure S3.**
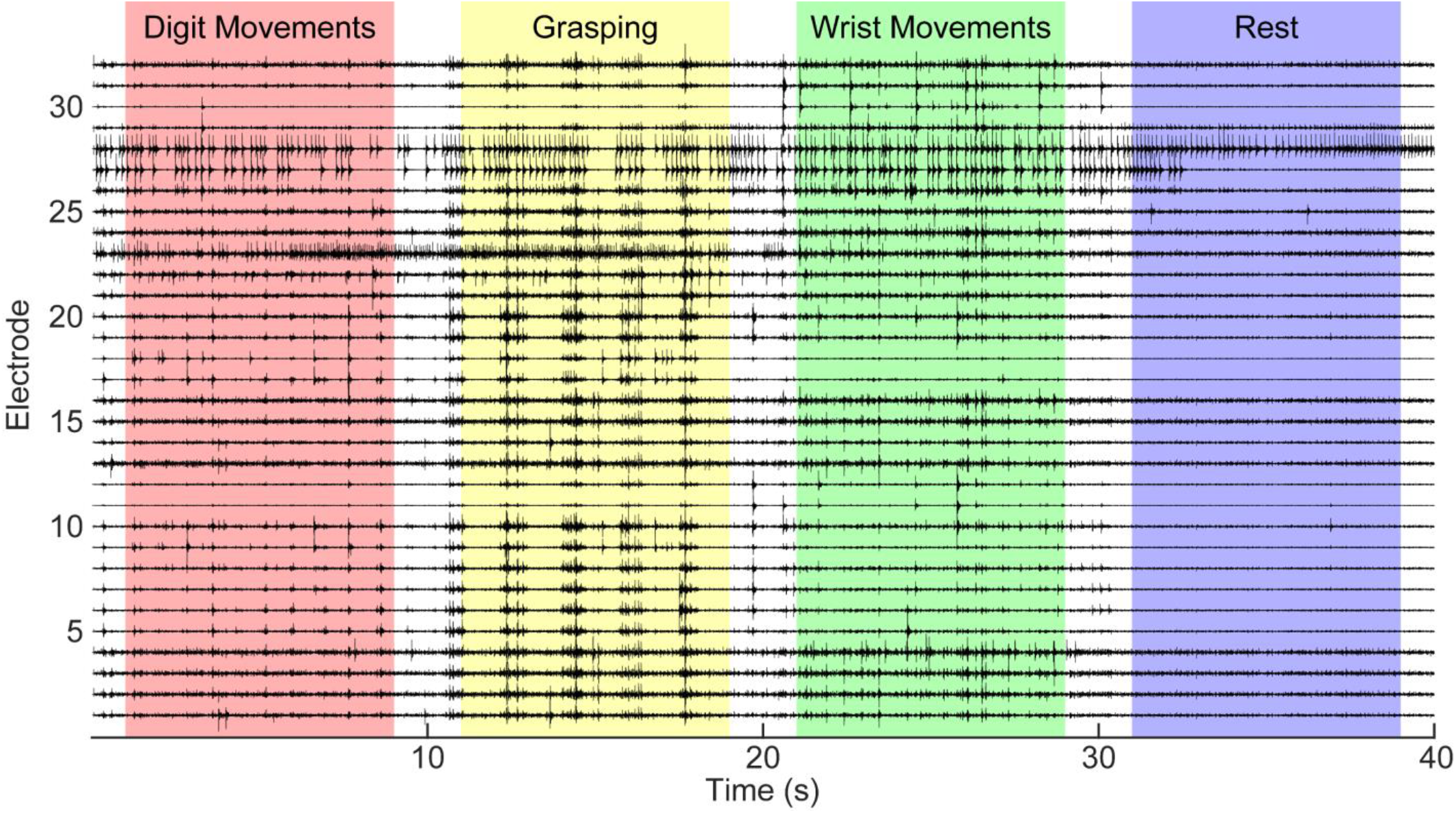
EMG activity recorded from iEMGs 502 days after they implanted in participant S7. EMG activity was still present for S7 despite long-term hand disuse and muscle atrophy prior to iEMG implantation. Traces show filtered and amplified EMG activity sampled at 1 kHz for all 32 electrodes. Aberrant tonic activity was present on some channels (e.g., e27 and e28), consistent with Complex Regional Pain Syndrome and myoclonus. Participants were instructed to perform individual digit movements (~0–10 seconds, red), grasping (~10–20 seconds, yellow), and wrist movements (~20–30 seconds, green), followed by a rest period (~30–40 seconds, violet).

**Figure S4.**
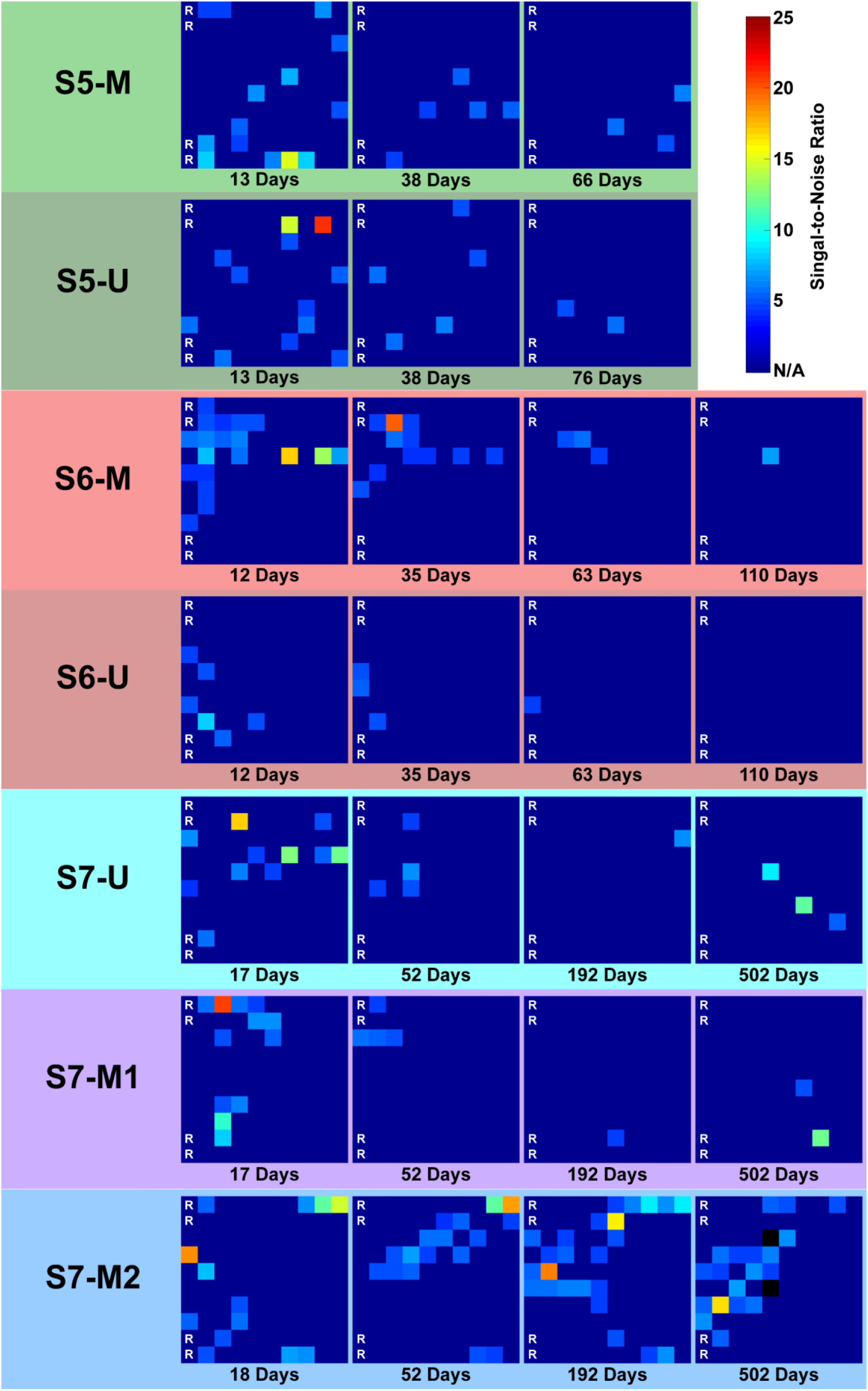
SNR of functional recording electrodes. Heatmaps show the 10×10 electrode grid on the USEA, with longest/deepest electrodes in column 1 on the left and shortest/shallowest electrodes in column 10 on the right. Electrodes 1, 11, 81, and 91 served as on-array references for recording. Hotter colors indicate a higher SNR. The darkest blue indicates the electrode was not functionally recording. Black color represents a SNR that is an outlier greater than 1.5 times the IQR.

**Figure S5.**
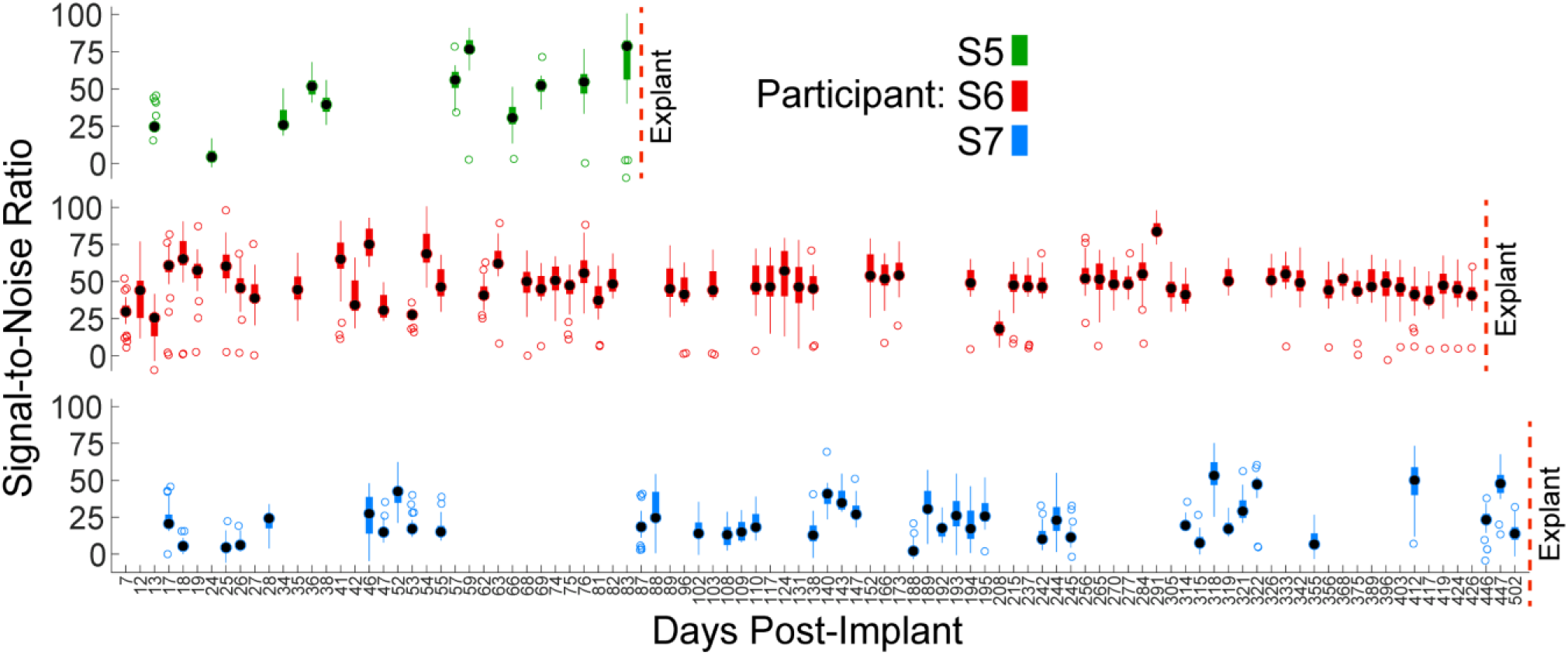
SNR of iEMG recordings over time. Timepoints are shown for each experimental session (ordinal scale). Black circles show median, thicker colored bars show IQR, thinner colored bars extend to the most extreme non-outlier values and open circles show outliers more than 1.5 times the IQR. Boxplots show the median SNRs of the 32 iEMG electrodes for each day, such that *N* = 32.

**Figure S6.**
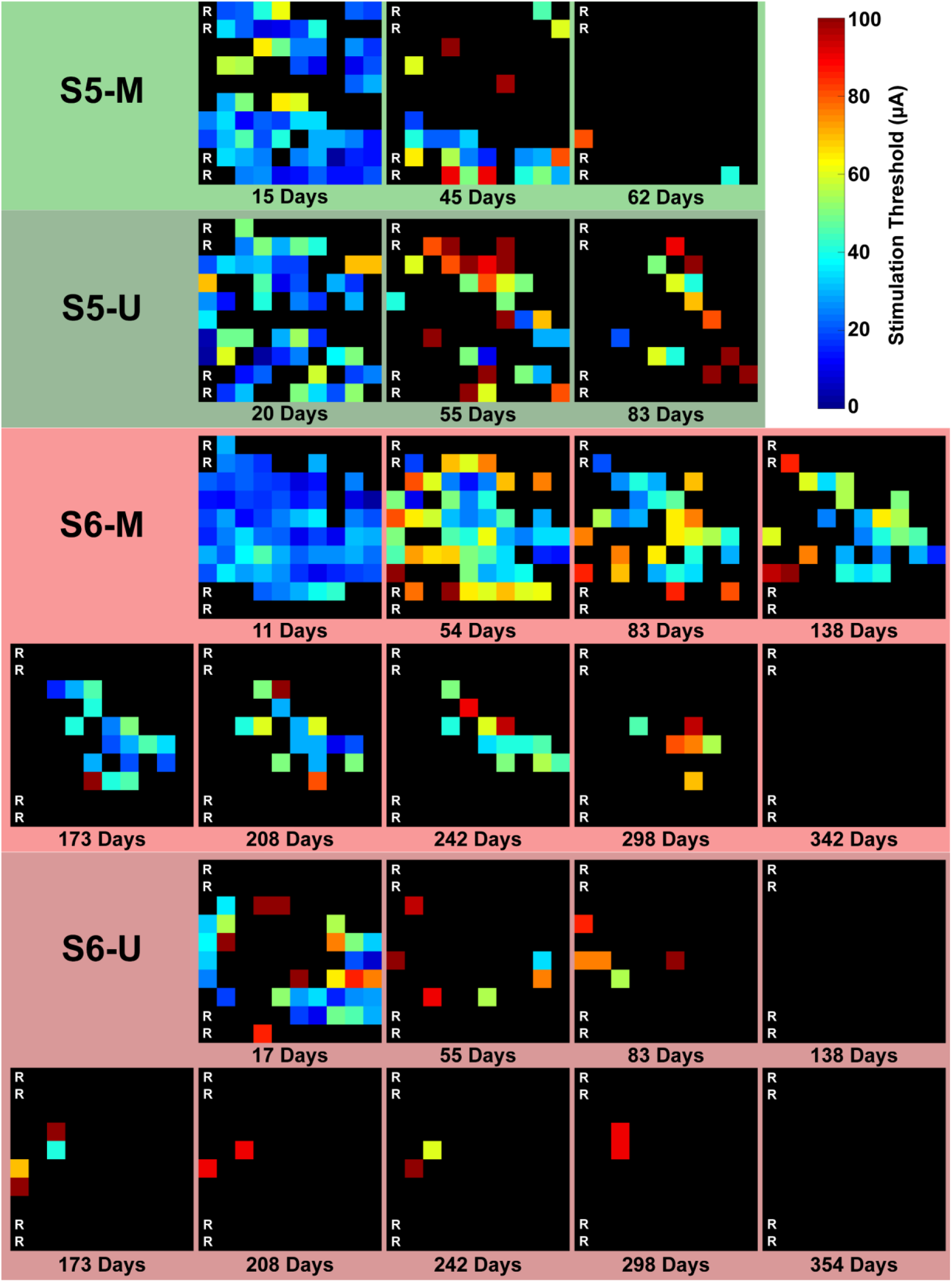
Detection threshold of USEA-evoked percepts over time. Heatmaps show the 10×10 electrode grid on the USEA, with longest/deepest electrodes in column 1 on the left and shortest/shallowest electrodes in column 10 on the right. Electrodes 1, 11, 81, and 91 served as on-array references for recording. Hotter colors indicate a higher detection threshold. Detection thresholds for each electrode were identified by increasing the stimulation amplitude in 1–10-μA step sizes until a percept was reliably detected.

